# Diagnostic Cerebrospinal Fluid Biomarker Discovery and Validation in Patients with Central Nervous System Infections

**DOI:** 10.1101/2020.01.13.899625

**Authors:** Tran Tan Thanh, Climent Casals-Pascual, Nguyen Thi Han Ny, Nghiem My Ngoc, Ronald Geskus, Le Nguyen Truc Nhu, Nguyen Thi Thu Hong, Du Trong Duc, Do Dang Anh Thu, Phan Nha Uyen, Vuong Bao Ngoc, Le Thi My Chau, Van Xuan Quynh, Nguyen Ho Hong Hanh, Nguyen Thuy Thuong Thuong, Le Thi Diem, Bui Thi Bich Hanh, Vu Thi Ty Hang, Pham Kieu Nguyet Oanh, Roman Fischer, Nguyen Hoan Phu, Ho Dang Trung Nghia, Nguyen Van Vinh Chau, Ngo Thi Hoa, Benedikt M. Kessler, Guy Thwaites, Le Van Tan

## Abstract

**Background:** Central nervous system (CNS) infections are common causes of morbidity and mortality worldwide. Rapid, accurate identification of the likely cause is essential for clinical management and the early initiation of antimicrobial therapy, which potentially improves clinical outcome.

**Methods:** We applied liquid chromatography tandem mass-spectrometry on 45 cerebrospinal fluid (CSF) samples from a cohort of adults with/without CNS infections to discover potential diagnostic protein biomarkers. We then validated the diagnostic performance of a selected biomarker candidate in an independent cohort of 364 consecutively treated adults with CNS infections admitted to a referral hospital in southern Vietnam.

**Results:** In the discovery cohort, we identified lipocalin 2 (LCN2) as a potential biomarker of bacterial meningitis. The analysis of the validation cohort showed that LCN2 could discriminate bacterial meningitis from other CNS infections, including tuberculous meningitis, cryptococcal meningitis and viral/antibody-mediated encephalitis (sensitivity: 0.88 (95% confident interval (CI): 0.77–0.94), specificity: 0.91 (95%CI: 0.88–0.94) and diagnostic odd ratio: 73.8 (95%CI: 31.8–171.4)). LCN2 outperformed other CSF markers (leukocytes, glucose, protein and lactate) commonly used in routine care worldwide. The combination of LCN2 and these four routine CSF markers resulted in the highest diagnostic performance for bacterial meningitis (area under receiver-operating-characteristic-curve 0.96; 95%CI: 0.93–0.99).

**Conclusions:** Our results suggest that LCN2 is a sensitive and specific biomarker for discriminating bacterial meningitis from a broad spectrum of CNS infections. A prospective study is needed to further assess the diagnostic utility of LCN2 in the diagnosis and management of CNS infections.

## INTRODUCTION

Central nervous system (CNS) infections cause significant mortality and morbidity worldwide, but especially in low- and middle-income countries (1). Common CNS infections include bacterial meningitis (BM), viral encephalitis, tuberculous meningitis (TBM) and cryptococcal meningitis (2), but there are >100 documented infectious causes of CNS infections (3). Additionally, over the last decade, antibody-mediated causes of encephalitis (e.g. anti-N-methyl-D-aspartate receptor (anti-NMDAR) encephalitis) have been recognized (4), which further challenges routine diagnostics.

Clinical features are often insufficient to discriminate the likely cause and standard laboratory investigations identify the causative agent in <60% of cases (5, 6). Critically, the clinical management of CNS infections varies according to its aetiology. Thus, rapid and accurate identification of the likely cause of the infection is essential to initiate appropriate therapy and improve patient outcome.

Over the last decade, mass-spectrometry has emerged as a sensitive, hypothesis-free approach for the discovery of novel diagnostic biomarkers in both communicable (e.g. CNS infections) and non-communicable diseases (7-9). However, previous biomarker-discovery studies of CNS infections have either been limited in sample size or have not included a validation phase (7). Here, using a mass-spectrometry based approach we first searched for novel diagnostic biomarkers in cerebrospinal fluid (CSF) samples from a discovery cohort of 45 patients with brain infections. We then sought to validate our findings in an independent cohort of 364 consecutively treated adults with CNS infections.

## MATERIALS AND METHODS

### Setting and the clinical studies

CSF samples were derived from three different clinical studies: study #1, #2 and #3 (Figure 1), conducted in the brain infections ward of the Hospital for Tropical Diseases (HTD) in Ho Chi Minh City, Vietnam. HTD is a tertiary referral hospital for severe infectious diseases, including suspected CNS infections, occurring in the southern provinces of Vietnam, with a population of >40 million.

**Figure 1:**
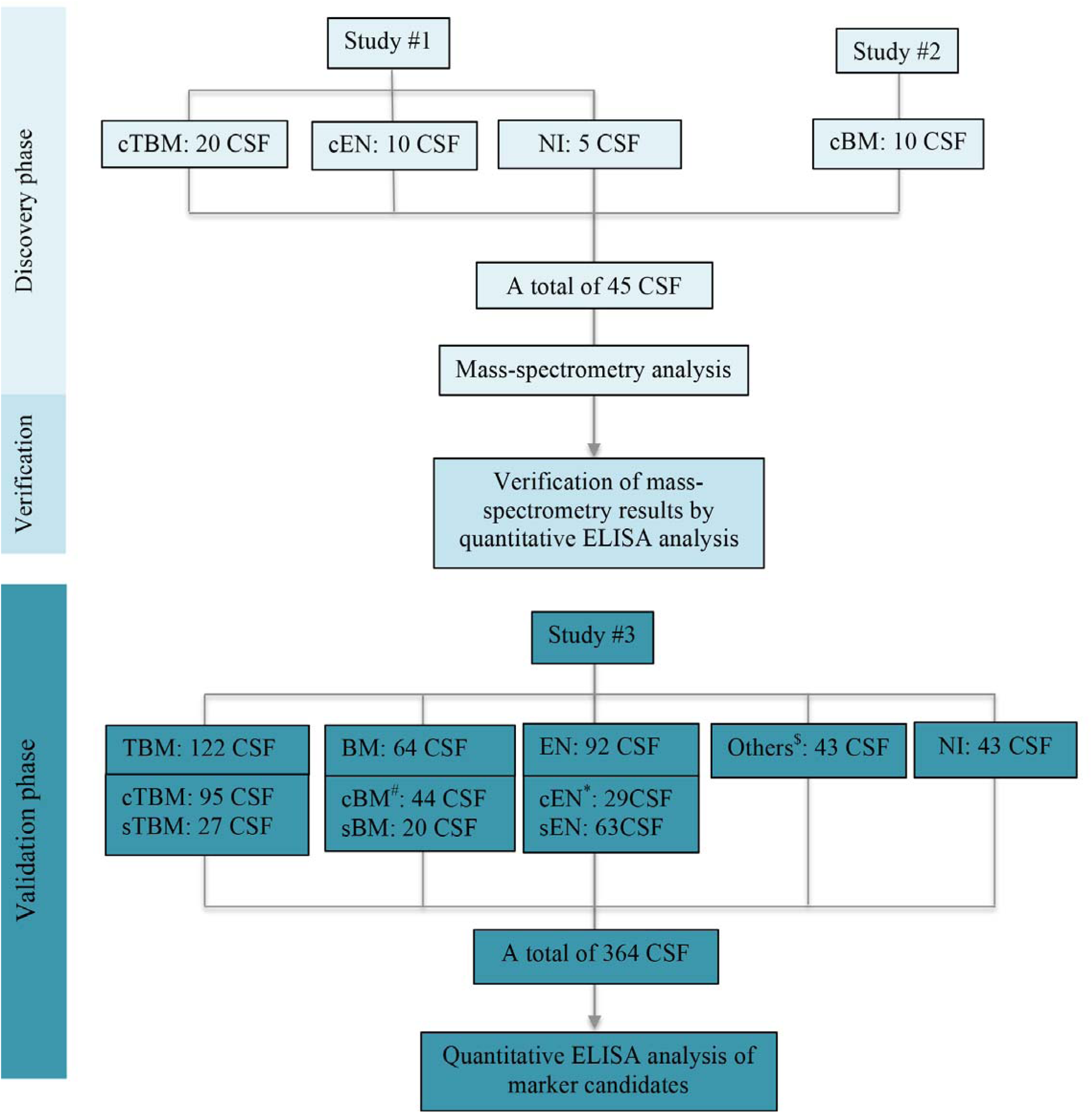
An overview of protein marker discovery phases and origin of clinical samples used for the analysis. **Note to Figure 1**: TBM: tuberculous meningitis, cTBM: confirmed tuberculous meningitis, sTBM: clinically suspected tuberculous meningitis, BM: bacterial meningitis, cBM: confirmed bacterial meningitis, sBM: clinically suspected bacterial meningitis, EN: encephalitis, cEN: confirmed encephalitis, sEN: clinically suspected encephalitis, NI: non-CNS infections ^#^Including: *S. suis* (n=20), *S. pneumonia* (n=6), *E. coli* (n=5), *N. meningitides* (n=2), *B. pseudomallei* (n=1), *E. faecalis* (n=1), *E. gallinarum* (n=1), *S. agalactiae* (n=1), *S. aureus* (n=1), *S. gallolyticus* (n=1) and gram staining positive only (n=5) ^*^including: herpes simplex virus (n=11), varicella zoster virus (n=7), dengue virus (n=5), Japanese encephalitis virus (n=2), dengue virus/Japanese encephalitis virus (n=1), mumps virus (n=1), measles virus (n=1) and influenza A virus (n=1) ^$^Including: cryptococcal meningitis (n=14), anti-NMDAR encephalitis (n=17), eosinophilic meningitis (n=10), neurotoxoplasmosis (n=2)

The clinical study #1 entitled “expanding the laboratory diagnosis of tuberculous meningitis and meningoencephalitis in Vietnam” was conducted during January 2015–September 2016 (10). As per the study protocol, any adult (≥18 years) with a suspected CNS infection and requirement for lumbar puncture was eligible for enrolment. Patients were excluded if pyogenic bacterial meningitis (very cloudy or pus-like CSF) was suspected, lumbar puncture was contra-indicated, or no informed consent was obtained.

The clinical study #2 focused on the immunological responses in bacterial meningitis patients, especially those infected with *Streptococcus suis*, and was conducted during 2015 and 2017. Any patient (≥16 years) with suspected pyogenic bacterial meningitis (very cloudy or pus-like CSF) was eligible for enrolment. Patient was excluded if lumbar puncture was contra-indicated, or no informed consent was obtained.

The clinical study #3 started in September 2017 and is on-going. The study aims to explore the potential diagnostic utility of next-generation sequencing and mass-spectrometry in CNS infections. Any patient (≥16 years) with suspected CNS infection and requirement for lumbar puncture was eligible for enrolment. Patients were excluded if no written informed consent was obtained.

For all the aforementioned studies, CSF and plasma samples were collected at presentation alongside demographic and clinical data and the results of routine diagnostic tests. All specimens were stored at −80°C until analysis.

### Routine diagnosis

As part of routine care, CSF specimens of patients with suspected CNS infections were cultured and/or examined by microscopy for the detection of bacteria, fungi and *Mycobacterium tuberculosis* with the use of standard methods (Table S1) (11). Herpes simplex virus (HSV) PCR was performed on CSF from those with suspected viral encephalitis. Varicella zoster virus (VZV) PCR, and serological testing for dengue virus (DENV) IgM, Japanese encephalitis virus (JEV) or mumps virus (MuV) was performed if clinically indicated and when testing for other pathogens was negative. Diagnosis of measles was based on compatible clinical features and the presence of measles IgM.

### Assignment of CNS infection diagnosis

Assignment of the CNS infection cause (TBM, BM, cryptococcal meningitis, eosinophilic meningitis, or anti-NMDAR encephalitis) was first based on the results of standard laboratory investigations. The diagnosis was confirmed if the relevant infectious agent was identified in the CSF. Otherwise, patients were considered as having clinically suspected CNS infections (TBM/BM/encephalitis) based on treatment responses and/or clinical judgment of treating physicians. Because of the focus of the present study, probable and possible TBM (defined by the Marais criteria (12)) were regarded as clinically suspected TBM. CNS infection was excluded in those with no meningeal signs, CSF laboratory parameters were in normal ranges, and all microbiological and serological investigations were negative.

### Sample preparation and mass-spectrometry analysis

CSF was analyzed as individual samples using proteomic platforms available at the Target Discovery Institute, University of Oxford. Briefly, CSF was digested in solution after reduction/alkylation with DTT/Iodoacetamide, followed by protein precipitation with Chloroform/Methanol (13). After tryptic digest, peptides were desalted using SepPak C18 cartridges (Waters) and injected into a LC-MS/MS platform consisting of Dionex Ultimate3000 nHPLC (Thermo) and either Q-Exactive or Q-Exactive HF. Peptides were separated with a linear gradient of 5-35% Acetonitrile in 0.1% Formic acid over 60 minutes using a 50cm x 75µm Easy Spray column (Thermo). MS1 resolution was set to 70,000 (HF 60,000) with an AGC target of 3E6. Fragment spectra were acquired with a resolution of 17,500 (HF 30,000) for up to 128ms (HF 45ms) and an AGC target of 1E5 ions. The output data were searched for human proteome (Uniprot, 05/2016 and 05/2018) using Mascot 2.5 or the Central Proteomics Pipeline (14). Peptide FDR was adjusted to 1% and lable free quantitation was conducted with either SINQ (15) within the CPFP or Progenesis QI version 3.1.4003.30577. Separation of CNS infection diagnostic groups based on the obtained peptide/protein profiles was performed using Perseus software version 1.6.6.0 (16)

### Measurement of lipocalin-2 concentration by quantitative ELISA

Measurement of lipocalin 2 (LCN2) concentrations was performed on CSF samples of the discovery and validation cohort as well as a subset of plasma samples of the validation cohort using Quantikine^®^ ELISA kits (R&D Systems, Minneapolis, MN, US). The experiments were performed according to the manufacturer’s instruction.

### Statistical analysis

Continuous variables were compared using the Mann-Whitney *U* test or the Kruskal-Wallis test or Wilcoxon signed-rank test. The correlation between continuous variables was assessed using Spearman correlation test. All statistical tests were performed two-sided. The area under the receiver operating characteristic curve (AUROC) was used to quantify the diagnostic performance of biomarkers for a given diagnosis. The cutoff values for outcome prediction were selected based on the highest sum of sensitivity and specificity. A logistic regression model was used to evaluate the diagnostic performance of two or more variables combined. All continuous variables were modeled as linear terms. All analyses were performed in SPSS V23.0 (IBM Corp, NY, US), and all figures were generated using GraphPad PRISM^®^ V5.04 (GraphPad Software Inc, CA, US).

### Ethics

The study was approved by Institutional Review Board of HTD and the Oxford Tropical Research Ethics Committee (OxTREC). Written informed consents were obtained from each participant or a relative if the patient was incapacitated.

## RESULTS

### Baseline characteristics of the study population

#### Discovery cohort

For the initial mass-spectrometry analysis, we selected a total of 45 patients enrolled in the clinical study #1 and #2. This consisted of 40 patients with laboratory confirmed CNS infections: TBM (n=20), BM (n=10), encephalitis (n=10), and five patients with non-CNS infection (Figure 1). Of the 10 patients with BM, seven were infected with *S. suis* and three with *S. pneumoniae*. Of the patients with encephalitis, herpes simplex virus was the cause in 5, DENV in 3, JEV in 1 and mumps virus in 1. The cohort’s clinical characteristics and outcomes are presented in Table S2.

#### Validation cohort

To validate the results of the discovery phase, we selected 364 consecutive adult patients enrolled in the study #3 (Figure 1). The baseline characteristics, clinical outcomes, and results of etiological investigations of the cohort are presented in Table S2 and the footnote of Figure 1, respectively. After the exclusion of 43 patients without CNS infections, the etiology was confirmed in 63% of the 321 patients with CNS infections. TBM was the most frequent diagnosis, followed by viral encephalitis and BM. The remaining patients included those with anti-NMDAR encephalitis, cryptococcal meningitis, parasitic eosinophilic meningitis and neurotoxoplasmosis (Figure 1). Of the patients with TBM, BM and viral encephalitis, a confirmed diagnosis was established in 97/122 (80%), 44/64 (69%) and 29/92 (32%), respectively. Of the 44 laboratory-confirmed bacterial meningitis patients, *S. suis*, was the commonest cause (n=20), followed by *S. pneumoniae* (n=6) and *Escherichia coli* (n=5). Of the 29 patients with laboratory confirmed viral encephalitis, HSV was the commonest cause (n=11), followed by VZV (n=7), and DENV (n=5) (Figure. 1).

### Biomarker discovery

Tandem mass-spectrometry analysis of 45 CSF samples of the discovery cohort identified a total of 1,012 proteins. Of these, 891 were included in the analysis based on the number of peptides and sequence coverage. Subsequent analysis identified a total of 729 quantifiable protein signatures that were clinical-entity specific, especially for patients with BM (Figure 2A). Of these, 60 and 19 were significantly expressed in the CSF of patients with BM and TBM, respectively (Table S3). No diagnostic biomarker candidate was found in patients with viral encephalitis.

**Figure 2.**
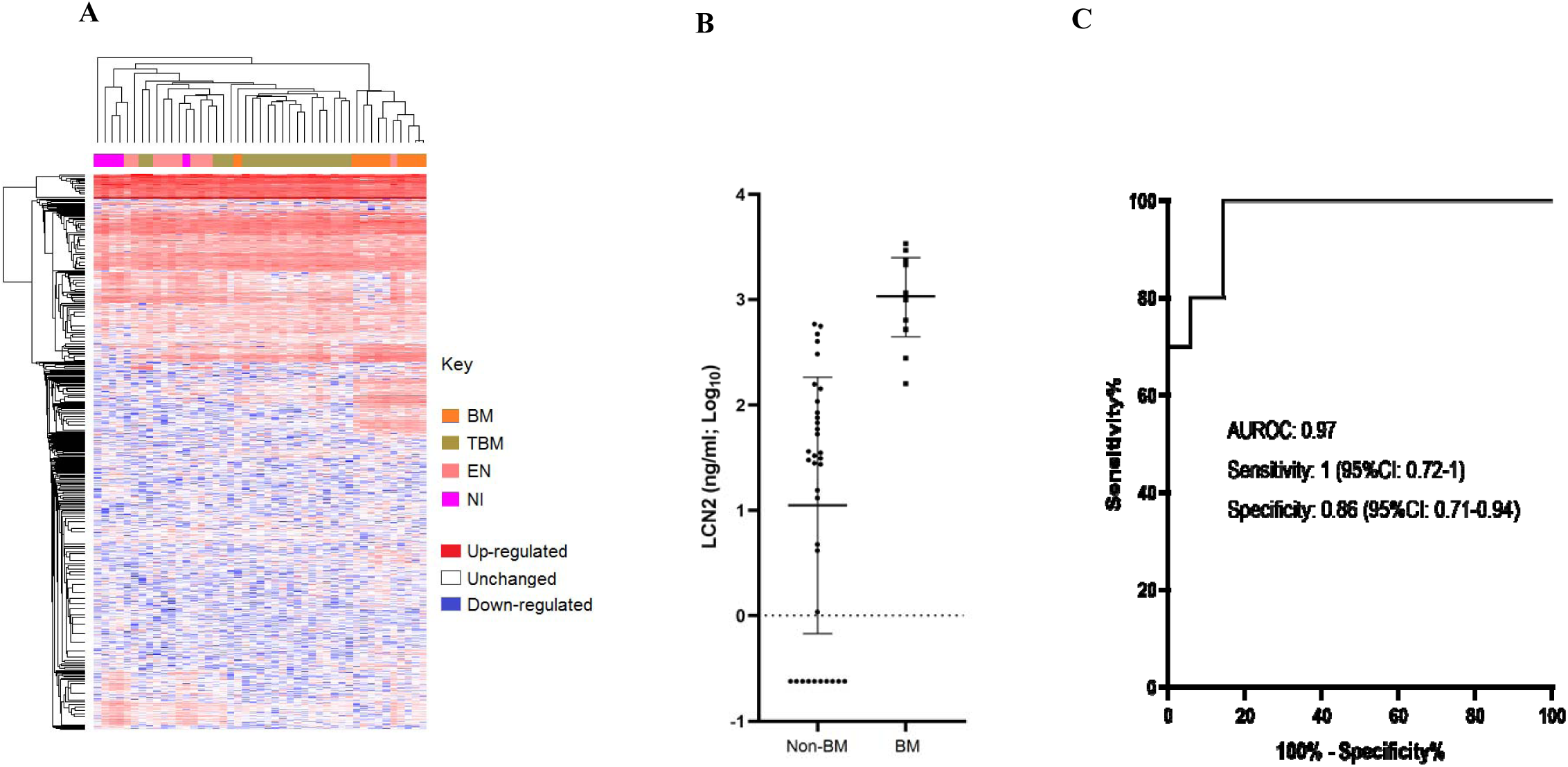
Results of mass-spectrometry and LCN2 ELISA analysis of the discovery cohort. (**A**) Heatmap showing clinical entities clustering based on the protein/peptide profiles obtained from label-free quantitative mass-spectrometry analysis of 45 patients of the discovery phase. Columns represent clinical entities, while rows represent individual proteins, (**B)** Dot plots demonstrating the difference in CSF LCN2 levels between BM and non-BM groups obtained from quantitative ELISA analysis, (**C**) AUROC curve based on LCN2 levels measured by quantitative ELISA analysis **Note to Figure 2**. Non-BM: non bacterial meningitis (encephalitis, tuberculous meningitis or non-CNS infections)

Of the protein candidates identified in the BM group, lipocalin 2 (LCN2), also known as neutrophil gelatinase-associated lipocalin, had a sensitivity of 1 (95%CI: 0.73-1) and a specificity of 0.89 (95%CI: 0.74-0.95) for prediction of BM (AUROC: 0.97 [95% confidence interval [CI], 0.9–1]). Because of its known biological significance in bacterial infections (17-19), previous reports of high concentrations in bacterial meningitis (20, 21), and the availability of a quantitative ELISA assay, LCN2 was thus selected for further evaluation.

### LCN2 ELISA analysis to verify the results of original LC-MS/MS analysis

In order to verify the mass-spectrometry findings, we performed quantitative ELISA analysis of the 45 CSF samples used for the discovery phase. Subsequently, the result suggested that LCN2 concentration of 159 ng/ml or above could accurately distinguish BM from TBM, encephalitis and non-CNS infections groups; AUROC curve: 0.97 (95%CI: 0.92–1), corresponding to the sensitivity of 1 (95%CI: 0.72–1) and the specificity of 0.86 (95%CI: 0.71–0.94) (Figures 2B and 2C). Thus, the diagnostic values of LCN2 based on the results of quantitative ELISA analysis confirmed the original finding of LC-MS/MS analysis.

### CSF LCN2 concentrations in the validation cohort

LCN2 was quantified in the CSF of the 364 consecutively treated adults with CNS infections enrolled in the study #3 (Figure 1). The results showed that LCN2 concentrations were significantly different amongst the diagnostic groups with the highest concentration observed in the BM group (median: 778. 8 ng/ml, range: 2.5–6566.3), followed by TBM groups (median: 86.3 ng/ml, range: 1.1–723.4) (Figure 3A). In contrast, LCN2 was almost absent or detected at very low levels in CSF of patients presenting with anti-NMDAR encephalitis (median: 0.9 ng/ml, range: 0.2–27.8) or in those without CNS infection (median: 0.2 ng/ml, range: 0.2–120.3) (Figure S1). Of the patients with BM, CSF LCN2 levels were higher in those with a confirmed diagnosis than in those without a bacteria identified (Figure S1), while the duration of illness at enrolment was similar between the two groups (data not shown).

**Figure 3.**
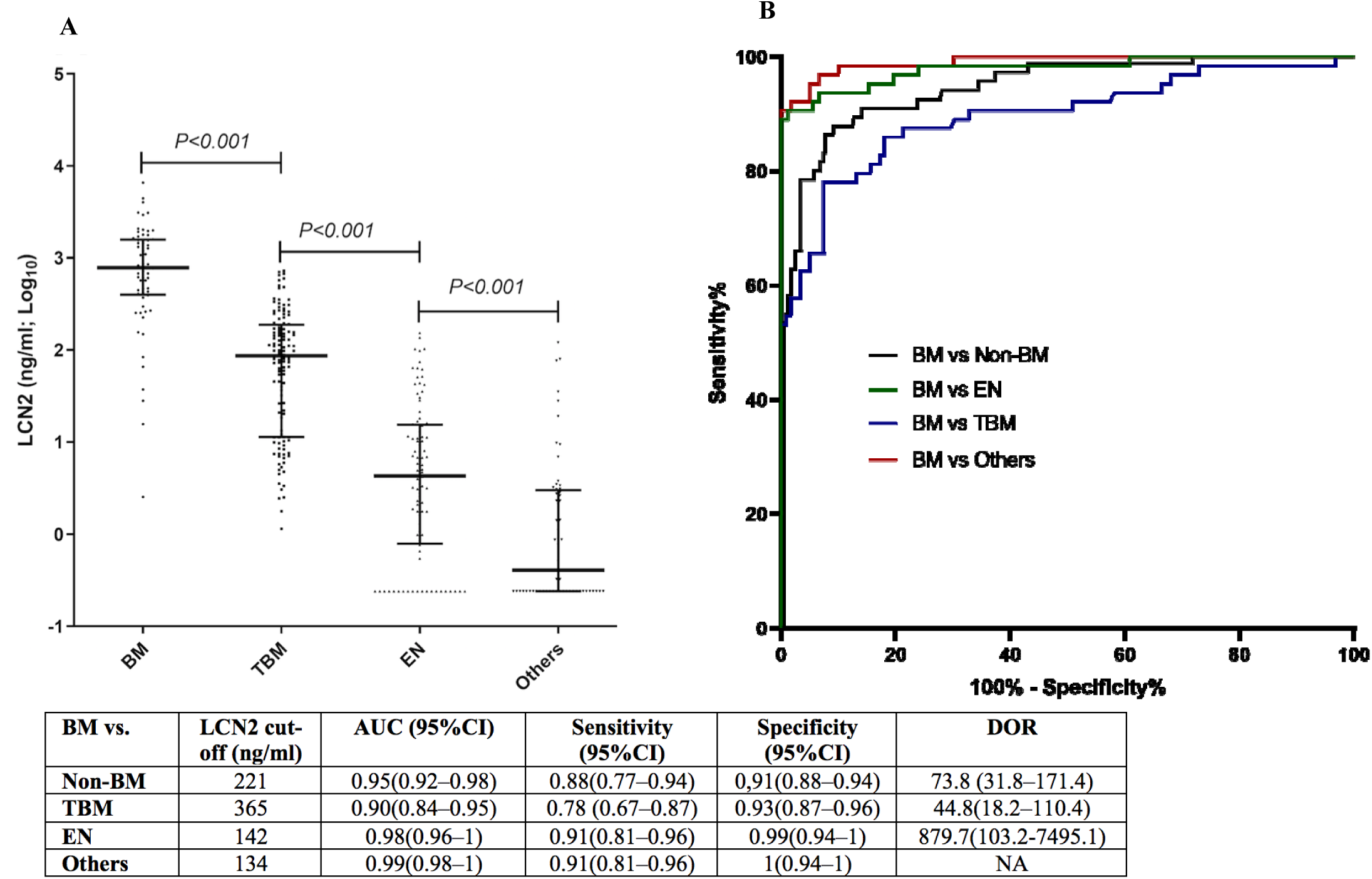
Results of LCN2 ELISA and AUROC analysis of the validation cohort. (**A**) LCN2 concentrations in patients with meningitis, tuberculous meningitis, encephalitis and others (cryptococcal meningitis, anti-NMDAR encephalitis, eosinophilic meningitis, neurotoxoplasmosis and non-CNS infections), (**B**) AUROC curves showing the diagnostic values of LCN2 in discriminating bacterial meningitis from other CNS infections entities **Note to Figure 3**: Others: patients with other CNS infections (cryptococcal meningitis, anti-NMDAR encephalitis, neurotoxoplasmosis, or eosinophilic meningitis) or non-CNS infections, Non-BM: non bacterial meningitis

### Diagnostic performance of CSF LCN2

Analysis of LCN2 concentrations obtained from the validation cohort demonstrated that LCN2 could accurately discriminate bacterial meningitis from other CNS infections with AUROC curves ranging from 0.9 (for BM vs. TBM, LCN2 concentration cut-off: 365 ng/ml) to 0.99 (BM vs. other CNS infections (i.e. non-encephalitis or non-TBM), LCN2 concentration cut-off: 134 ng/ml) and a diagnostic odd ratio (DOR) of 44.8 or above (Figure 3B).

Currently, CSF parameters such as leukocytes, protein, lactate and glucose concentrations are routinely used as diagnostic makers in the primary assessment of patients presenting with CNS infections. We thus compared the diagnostic performance of LCN2 alone and in combination with these markers..

LCN2 outperformed the existing biomarkers in discriminating between BM and other CNS infections (including TBM and encephalitis) (Figures 4A and 4B). When LCN2 was combined with leukocytes, protein, lactate and glucose concentrations in CSF, the diagnostic model consisting of LCN2 and these four CSF parameters provided the highest discriminatory ability for BM (Figure 4C). More specifically, in terms of discriminating between BM and all other CNS infections, the predictive values for BM based on AUROC curves and DOR increased from 0.94 (95%CI: 0.80–0.98) to 0.96 (95%CI: 0.93–0.99), and 66.2 to 308.3 when LCN2 was added to the CSF parameters based model (Figure 4C). Similar results were obtained when assessing the utility of LCN2 in discriminating between BM and other specific clinical entities (TBM or encephalitis) (Figure 4D and Figure S2). LCN2 did not, however, help distinguish confirmed from suspected BM (Table S4).

**Figure 4.**
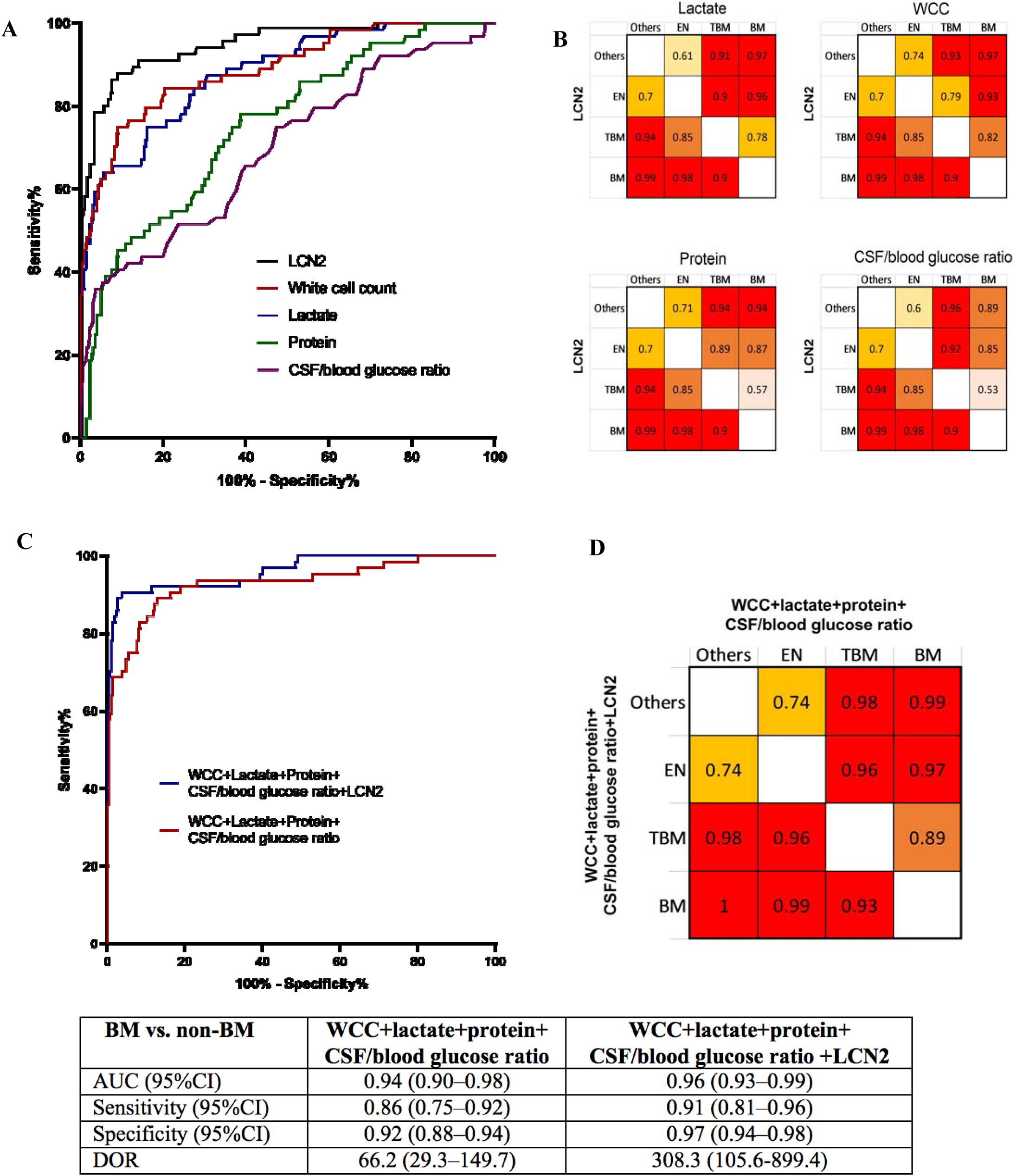
Diagnostic values of LCN2 in predicting bacterial meningitis in comparison or in combination with existing CSF parameters. (**A**) AUROC curves showing that LCN2 is better than the existing CSF parameters in distinguishing between bacterial meningitis with other CNS infections (tuberculous meningitis, encephalitis, anti-NMDAR encephalitis, crytococcal meningitis, neurotoxoplasmosis, eosinophilic meningitis or non-CNS infections), (**B**) AUCROC values of subgroup analyses, (**C**) AUROC curves showing that LCN2 significantly improves the discriminatory ability of the diagnostic model for bacterial meningitis using the remaining CNS infections groups as controls, (**D**) AUROC values of subgroup analyses **Note to Figure 4:** WCC: white blood cell count (leukocyte count)

### Association between CSF and plasma concentrations of LCN2

LCN2 is a ubiquitous protein which can be found in bodily fluids of healthy individuals (22). We assessed if plasma LCN2 can be a surrogate of CSF LCN2 in a subset of 22 patients with BM (laboratory confirmed: n=14 and clinically suspected: n=8). Plasma LCN2 concentration was, however, significantly lower than that of CSF; median: 147.9 ng/ml, range: 33.7–194.8 vs. CSF LCN2: median: 472.1 ng/ml, range: 15.7–3102.3, P<0.001. There was no correlation between CSF and plasma LCN2 (Spearman R: 0.37, P=0.08), suggesting that LCN2 is intrathecally produced in response to the bacterial invasion of the CNS.

## DISCUSSION

Here, using a mass-spectrometry-based approach, we initially identified LCN2 as a potential diagnostic marker for BM. Additional validation work on an independent cohort showed that LCN2 could accurately discriminate BM from other CNS infections. LCN2 also outperformed existing BM diagnostic makers (CSF leukocytes, and protein, glucose and lactate concentrations) that are currently used as part of routine care. A diagnostic model consisting of LCN2 and these four CSF parameters gave the best diagnostic performance for BM. Our data thus suggest that LCN2 can act as an independent diagnostic maker of BM alone or in combination with other CSF parameters.

LCN2 is secreted by neutrophils, hepatocytes and renal tubular cells (23). It is encoded by LCN2 gene and is known to have antibacterial properties because of its ability to inhibit the bacterial growth via the interference of bacterial iron uptake (23). LCN2 has recently been recognized as a sensitive biomarker for the diagnosis of severe blood stream infection (24) and pneumonia caused by *S. pneumoniae* (18). High concentrations of LCN2 in the CSF of patients with BM have been previously reported (20, 21). However, previous studies only focused on quantifying LCN2 concentrations in patients with confirmed BM and viral encephalitis and did not compare the performance of LCN2 against commonly used CSF markers such as leukocytes, glucose, protein and lactate. Our study was conducted in Vietnam and included patients with many different CNS infections, including bacterial, fungal, tuberculous, viral and parasitic meningitis, and anti-NMDAR encephalitis). Additionally, we also compared the diagnostic performance of LCN2 against that of CSF markers commonly used as part of routine care worldwide. As such, our results have expanded our knowledge about the relation between LCN2 and CNS infections, and for the first time provide strong evidence that LCN2 is a highly sensitive biomarker for discriminating BM from a broad-spectrum of CNS infections.

The differences in CSF LCN2 levels between laboratory confirmed and clinically suspected BM groups pointed to the association between the host responses and an on-going infection (i.e. the presence of a bacterial pathogen in clinical samples at the time of collection). This is in agreement with previous studies showing that the decrease of plasma LCN2 level was correlated with the success of antibiotic treatment in patients with bacteremia (17). Collectively, CSF LCN2 might also be a useful marker for treatment response assessment. Therefore, further research should aim at defining the optimal cut-off of LCN2 concentrations that can be used to inform the administration or withdrawal of antibiotics in patients with BM.

Our study has some of limitations. A part from LCN2, we did not explore the utility potential of the other biomarker candidates identified in the discovery cohort (e.g. CSF cathelicidin for BM (25)) detected by original mass-spectrometry analysis). Likewise, we did not assess the diagnostic performance of LCN2 against and/or in combination newly proposed biomarkers for CNS infections such as procalcitonin and heparin-binding protein for BM (7, 9, 26-28), and CSF lipoarabinomannan for TBM (29). Additionally, we only focused our analysis on adults, leaving the utility potential of LCN2 in pediatric CNS infections unknown.

In spite of these limitations, the strengths of our study include that it represents the largest and most comprehensive mass-spectrometry-based biomarker discovery investigation focusing on patients with various clinical entities of CNS infections to date (7). The study was also conducted in Vietnam and therefore includes all the major infectious causes of CNS infections seen globally. Additionally, because our study was conducted at a single major tertiary referral hospital, all routine diagnostic approaches and patient assessments were consistent over the course of the study, thereby minimizing potential bias.

To summarize, our study showed for the first time that LCN2 is a highly sensitive biomarker for accurate prediction of BM in adults, especially when used alongside other standard CSF parameters. Prospective studies are needed to assess the utility potential of LCN2 in the diagnosis and management of CNS infections, including children, and whether it can be used in settings with limited laboratory capacity to improve outcomes from these devastating conditions.

## ACKNOWLEDGEMENTS

We thank Le Kim Thanh, Pham Thi Kim, Vu Thi Mong Dung and Lam Anh Nguyet for their logistic support, and Dr Nguyen Thi Hoang Mai for her help with material collection. We are indebted to patients for their participations in this study and the doctors and nurses of Viet Anh ward, HTD, who cared for the patients.

This study was funded by the Wellcome Trust of Great Britain (106680/B/14/Z and 204904/Z/16/Z awarded to GT and LVT, respectively).

## SUPPLEMENTARY MATERIALS

**Table S1.** Diagnostic tests carried out as part of routine care and/or as per the study protocols.

|  | **Study #1** | **Study #2** | **Study #3** |
| --- | --- | --- | --- |
| Gram stain | Y | Y | Y |
| Bacterial culture | Y | Y | Y |
| Ziehl-Neelsen staining | Y | Y | Y |
| GenXpert | Y | Y | Y |
| MGIT | Y | Y | Y |
| *S. suis* PCR | Y | Y | Y |
| *S. pneumoniae* PCR | Y | Y | Y |
| *N. meningitidis* PCR | Y | Y | Y |
| 16S rRNA PCR | ND | Y | Y |
| HSV PCR | Y | Y | Y |
| VZV PCR | Y | Y | Y |
| DENV PCR | ND | ND | Y |
| JEV PCR | ND | ND | Y |
| Flavivirus PCR | ND | ND | Y |
| Enterovirus PCR | ND | ND | Y |
| Influenza A virus | Y | Y | Y |
| Mumps virus PCR | ND | ND | Y |
| Zika virus PCR | ND | ND | Y |
| *Angiostrongylus cantonensis* PCR | ND | ND | Y |
| Cryptococcal FLA | Y | Y | Y |
| DENV serology | Y | Y | Y |
| JEV serology | Y | Y | Y |
| Anti-NMDAR | ND | Y | Y |

**Note to Table S1:** Y: yes. ND: not done

**Table S2.** Baseline characteristics of the discovery and validation cohort.

|  | Discovery cohort |  |  |  | Validation cohort |  |  |  |  |  |  |
| --- | --- | --- | --- | --- | --- | --- | --- | --- | --- | --- | --- |
|  | TBM (N=20) | Encephalitis (N=10) | BM<sup>*</sup> (N=10) | Non-CNS infections (N=5) | BM (N=64)<sup>#</sup> | TBM (N=122)<sup>‡</sup> | Encephalitis (N=92)<sup>§</sup> | Anti-NMDAR (N=17) | Eosinophilic meningitis (N=10) | Cryptococcal meningitis (N=14) | Non-CNS infections (N=43) |
| **Demographics** |  |  |  |  |  |  |  |  |  |  |  |
| Age in years | 40(23-75) | 33(18-53) | 49(23-74) | 58(0-70) | 55 (17-87) | 41(17-87) | 31 (16-78) | 25(17-48) | 30(18-60) | 35.5(22-68) | 48(20-92) |
| Gender | 11/9 | 8/2 | 7/3 | 2/3 | 44/20 | 97/25 | 54/38 | 9/8 | 4/6 | 10/4 | 25/18 |
| Ho Chi Minh City origin | 5(25) | 2(20) | 4(44.4) | 2(40) | 12(18.8) | 29(23.8) | 27(29.3) | 3(17.6) | 2(20) | 4(28.6) | 10(23.3) |
| **Illness day at enrollment** | 14(0-60) | 5(0-10) | 2(0-13) | 4(3-18) | 4 (1-30) | 12 (2-90) | 6(1-90) | 22(6-37) | 25.5(7-60) | 18(4-30) | 5(1-60) |
| **Length of hospital stay** | 24(0-59) | 5(0-67) | 18(13-26) | 2(1-17) | 14 (1-119) | 26(0-162) | 11(0-118) | 41(27-102) | 11(1-23) | 23(0-134) | 8(0-75) |
| **Clinical signs/symptoms** |  |  |  |  |  |  |  |  |  |  |  |
| Fever (n,%) | 19(95) | 10(100) | NA | 5(100) | 59(96.7) | 117(96.7) | 83(92.2) | 13(76.5) | 7(70) | 12(85.7) | 36(87.8) |
| Headache (n,%) | 18(90) | 10(100) | 6(66.7) | 3(60) | 57(91.9) | 113(96.6) | 64(76.2) | 9(64.3) | 10(100) | 13(92.9) | 15(40.5) |
| Cranial nerve palsy (n,%) | 4(20) | 2(20) | NA | 0 | 5(7.8) | 23(18.9) | 9(9.8) | 0 | 3(30) | 4(28.6) | 3(7) |
| Hemiplegia (n,%) | 1(5) | 2(20) | NA | 0 | 1(1.6) | 12(9.8) | 4(4.3) | 0 | 1(10) | 1(7.1) | 3(7) |
| Paraplegia (n,%) | 0 | 0 | NA | 0 | 1(1.6) | 10(8.2) | 3(3.3) | 0 | 1(10) | 1(7.1) | 4(9.3) |
| Tetraplegia (n,%) | 0 | 0 | NA | 0 | 1(1.6) | 5(4.1) | 3(3.3) | 0 | 0 | 1(7.1) | 4(9.3) |
| Convulsions (n,%) | 1(5) | 1(10) | NA | 0 | 1(1.6) | 2(1.7) | 15(16.5) | 0 | 0 | 0 | 5(11.9) |
| Neck stiffness (n,%) | 17(85) | 9(90) | NA | 4(80) | 47 (77) | 57(47.6) | 30(33.7) | 5(29.4) | 5(50) | 7(50) | 8(19.5) |
| GCS** at enrolment (median, range) | 14(8-15) | 11(7-15) | NA | 14(10-15) | 12 (3-15) | 14(4-15) | 11(3-15) | 11(6-14) | 15(9-15) | 13(8-15) | 13(7-15) |
| HIV positive, (n%) | 2(20) | 1(10) | NA | 0 | 0 | 22(24.2) | 0 | 0 |  |  | 1(2.3) |
| **CSF examinations** |  |  |  |  |  |  |  |  |  |  |  |
| CSF leukocyte count (per mm3) | 317 (58-896) | 209 (18-1571) | 11000 (500-19200) | 5(1-93) | 1924 (24-51810) | 312(3-3969) | 43(1-909) | 23(6-187) | 501(140-1101) | 36.5(2-357) | 2(1-2700) |
| CSF neutrophils (%) | 26(3-93) | 3(0-18) | 91.5(83-98) | 20(0-87) | 83 (10-98) | 26(0-95) | 14(0-91) | 14(9-65) | 12(7-42) | 26.5(0-67) | 30(0-93) |
| CSF lymphocytes (%) | 74(7-94) | 96(0-98) | 6(0.9-17) | 13(0-99) | 17 (2-90) | 73(5-92) | 84(0-94) | 86(35-91) | 47(20-70) | 67.5(33-86) | 50(0-99) |
| CSF/blood glucose ratio | 0.19<br>(0.04-0.4) | 0.61<br>(0.5-7.66) | NA | 0.65<br>(0.41-0.67) | 0.3 (0-1) | 0.3(0.1-0.7) | 4(1.7-7.7) | 0.8(0.5-1.4) | 0.4(0.4-0.9) | 0.3(0-0.6) | 0.7(0.4-1.3) |
| CSF lactate (mmol/L) | 6.77<br>(3.21-12.43) | 2.79<br>(1.88-3.52) | NA | 3.65<br>(1.78-7.3) | 9.9 (2.3-28.2) | 5(1.9-12.8) | 2.5(1.3-6.6) | 1.9(1.4-2.7) | 2.7(2.2-4.2) | 4.9(2.7-12.7) | 2.5(1.4-6.4) |
| Total protein (g/L) | 2(1.1-4.1) | 0.8(0.3-2.4) | NA | 0.5(0.3-2.6) | 2.3 (0.3-8.7) | 1.9(0.2-29.8) | 0.7(0.1-3.2) | 0.3(0.2-0.8) | 0.8(0.2-3.9) | 0.6(0.4-1.8) | 0.4(0.2-3.2) |
| **Discharge mRS<sup>^</sup>** |  |  |  |  |  |  |  |  |  |  |  |
| 0 | 3(15) | 0 | NA | 1(20) | 8(12.5) | 22(18) | 17(18.2) | 0 | 0 | 0 | 5(11.6) |
| 1 | 3(15) | 2(20) | NA | 0 | 6(9.4) | 16(13.1) | 19(20.7) | 2(11.8) | 3(30) | 0 | 6(14) |
| 2 | 5(25) | 0 | NA | 2(40) | 12(18.8) | 15(13.2) | 19(20.7) | 0 | 4(40) | 1(7.1) | 8(18.6) |
| 3 | 4(20) | 4(40) | NA | 1(20) | 20(31.3) | 14(11.5) | 12(13) | 6(35.3) | 1(10) | 4(28.6) | 8(18.6) |
| 4 | 1(5) | 2(20) | NA | 0 | 11(17.2) | 22(18) | 13(14.1) | 3(17.6) | 1(10) | 3(21.4) | 9(20.9) |
| 5 | 2(10) | 2(20) | NA | 0 | 3(4.7) | 17(13.7) | 10(10.9) | 5(29.4) | 1(10) | 1(7.1) | 6(14) |
| 6 | 2(10) | 0 | NA | 1(20) | 4(6.3) | 16(13.1) | 2(2.2) | 1(5.9) | 0 | 5(35.7) | 1(2.3) |

**Note to Table S2:** <sup>†</sup>outcomes at discharge were recorded as full recovery (n=4) or neurological deficit (n=4)\*due to the small sample size, data on two cases with neurotoxoplasmosis are not shown, <sup>#</sup>including 44 laboratory confirmed cases, <sup>‡</sup>including 95 laboratory confirmed cases, <sup>§</sup>including 23 laboratory confirmed cases, \*\*Glasgow coma score, <sup>^</sup>Modified Rankin Scale (0: full recovery with no symptoms, 1: No significant disability, 2: Slight disability, 3: Moderate disability, 4: Moderately severe disability, 5: Severe disability, and 6: Dead); BM: bacterial meningitis, TBM: tuberculous meningitis; Data are number (%), continuous variables are presented as median (range)

**Table S3.** List of marker candidates identified by mass spectrometry analysis.

| BM group |  |  |  |  |  |  |  |
| --- | --- | --- | --- | --- | --- | --- | --- |
| No | Protein ID | Protein name | Gene name | Mean intensity of BM (log<sub>2</sub>) | Mean intensity of Other (log<sub>2</sub>) | Difference in intensity between BM and Others | -Log (p value) |
| 1 | P06744 | Glucose-6-phosphate isomerase | GPI | -19.99 | -24.54 | -4.55 | 7.74 |
| 2 | P60660-2 | Myosin light polypeptide 6 | MYL6 | -19.77 | -26.05 | -6.28 | 7.3 |
| 3 | P28676 | Grancalcin | GCA | -21.18 | -26.58 | -5.4 | 7.11 |
| 4 | P11413-2 | Glucose-6-phosphate 1-dehydrogenase | G6PD | -21.17 | -26.4 | -5.23 | 6.31 |
| 5 | P26583 | High mobility group protein B2 | HMGB2 | -20.83 | -26.23 | -5.41 | 5.3 |
| 6 | P05109 | Protein S100-A8 | S100A8 | -14.87 | -20.83 | -5.96 | 5.87 |
| 7 | P05164-2 | Myeloperoxidase | MPO | -18.03 | -23.71 | -5.68 | 5.84 |
| 8 | P06702 | Protein S100-A9 | S100A9 | -14.55 | -21.29 | -6.75 | 5.84 |
| 9 | P43490 | Nicotinamide phosphoribosyltransferase | NAMPT | -21.54 | -26.07 | -4.53 | 5.82 |
| 10 | P80188-2 | Neutrophil gelatinase-associated lipocalin | LCN2 | -17.82 | -23.6 | -5.78 | 5.81 |
| 11 | P22894 | Neutrophil collagenase | MMP8 | -19.34 | -24.07 | -4.75 | 5.77 |
| 12 | P50395 | Rab GDP dissociation inhibitor beta | GDI2 | -20.2 | -24.66 | -4.46 | 5.74 |
| 13 | P20160 | Azurocidin | AZU1 | -20.45 | -26.12 | -5.67 | 5.61 |
| 14 | P41218 | Myeloid cell nuclear differentiation antigen | MNDA | -19.89 | -24.52 | -4.63 | 5.60 |
| 15 | P61160 | Actin-related protein 2 | ACTR2 | -21.19 | -25.46 | -4.27 | 5.47 |
| 16 | O15144 | Actin-related protein 2/3 complex subunit 2 | ARPC2 | -19.86 | -25.15 | -5.29 | 5.47 |
| 17 | P08670 | Vimentin | VIM | -18.54 | -22.2 | -3.66 | 5.46 |
| 18 | P08107 | Heat shock 70 kDa protein 1A | HSPA1A | -19.45 | -24.1 | -4.65 | 5.37 |
| 19 | P30044-2 | Peroxisredoxin-5, mitochondrial | PRDX5 | -20.84 | -25.66 | -4.82 | 5.23 |
| 20 | P04040 | Catalase | CAT | -19.23 | -24.71 | -5.48 | 5.22 |
| 21 | P09429 | High mobility group protein B1 | HMGB1 | -21.35 | -25.89 | -4.53 | 5.12 |
| 22 | P61158 | Actin-related protein 3 | ACTR3 | -20.73 | -25.39 | -4.66 | 5.03 |
| 23 | P35579 | Myosin-9 | MYH9 | -21.18 | -26.72 | -5.54 | 5.02 |
| 24 | P04083 | Annexin A1 | ANXA1 | -19.86 | -25.38 | -5.52 | 4.81 |
| 25 | P49913 | Cathelicidin antimicrobial peptide | CAMP | -20.72 | -24.6 | -3.88 | 4.74 |
| 26 | P12814-3 | Alpha-actinin-1 | ACTN1 | -20.87 | -25.06 | -4.18 | 4.73 |
| 27 | U3KPS2 | Myeloblastin | PRTN3 | -19.02 | -22.98 | -3.96 | 4.71 |
| 28 | P01040 | Cystatin-A | CSTA | -18.79 | -24.16 | -5.37 | 4.7 |
| 29 | Q6UX06 | Olfactomedin-4 | OLFM4 | -22.21 | -26.55 | -4.33 | 4.69 |
| 30 | P52209-2 | 6-phosphogluconate dehydrogenase, decarboxylating | PGD | -19.4 | -24.03 | -4.63 | 4.67 |
| 31 | P37837 | Transaldolase | TALDO1 | -19.90 | -24.88 | -4.98 | 4.6 |
| 32 | P51149 | Ras-related protein Rab-7a | RAB7A | -21.76 | -25.96 | -4.21 | 4.59 |
| 33 | P08246 | Neutrophil elastase | ELANE | -17.89 | -23.13 | -5.24 | 4.59 |
| 34 | O15143 | Actin-related protein 2/3 complex subunit 1B | ARPC1B | -21.64 | -26.41 | -4.77 | 4.58 |
| 35 | O43707 | Alpha-actinin-4 | ACTN4 | -21.84 | -25.85 | -4.01 | 4.52 |
| 36 | P08311 | Cathepsin G | CTSG | -19.38 | -24.24 | -4.86 | 4.48 |
| 37 | P59998-3 | Actin-related protein 2/3 complex subunit 4 | ARPC4 | -19.92 | -24.16 | -4.24 | 4.39 |
| 38 | P61626 | Lysozyme C | LYZ | -16.34 | -18.05 | -1.71 | 4.39 |
| 39 | P30041 | Peroxisredoxin-6 | PRDX6 | -21.56 | -25.35 | -3.79 | 4.35 |
| 40 | P00338-3 | L-lactate dehydrogenase A chain | LDHA | -20.19 | -23.1 | -2.91 | 4.24 |
| 41 | Q05315 | Galectin-10 | CLC | -21.36 | -25.34 | -3.98 | 4.18 |
| 42 | P09960 | Leukotriene A-4 hydrolase | LTA4H | -21.47 | -24.95 | -3.48 | 4.15 |
| 43 | O14950 | Myosin regulatory light chain 12B | MYL12B | -21.26 | -25.66 | -4.4 | 4.12 |
| 44 | P09211 | Glutathione S-transferase P | GSTP1 | -18.82 | -23.38 | -4.56 | 4.1 |
| 45 | P00491 | Purine nucleoside phosphorylase | PNP | -21.14 | -25.57 | -4.43 | 4.07 |

| 46 | P18428 | Lipopolysaccharide-binding protein | LBP | -20.82 | -24.91 | -4.09 | 4.05 |
| --- | --- | --- | --- | --- | --- | --- | --- |
| 47 | P60709 | Actin, cytoplasmic 1 | ACTB | -16.18 | -17.82 | -1.65 | 4.02 |
| 48 | P21333-2 | Filamin-A | FLNA | -21.85 | -26.23 | -4.38 | 4.01 |
| 49 | Q9ULZ3-2 | Apoptosis-associated speck-like protein containing a CARD | PYCARD | -20.63 | -24.76 | -4.13 | 3.88 |
| 50 | P47756-2 | F-actin-capping protein subunit beta | CAPZB | -21.9 | -26.33 | -4.44 | 3.85 |
| 51 | P62491-2 | Ras-related protein Rab-11A | RAB11A | -21.91 | -26.14 | -4.23 | 3.82 |
| 52 | Q01518 | Adenylyl cyclase-associated protein 1 | CAP1 | -20.93 | -25.4 | -4.47 | 3.77 |
| 53 | O15145 | Actin-related protein 2/3 complex subunit 3 | ARPC3 | -21.08 | -24.75 | -3.67 | 3.77 |
| 54 | O00299 | Chloride intracellular channel protein 1 | CLIC1 | -21.69 | -26.07 | -4.38 | 3.75 |
| 55 | P35754 | Glutaredoxin-1 | GLRX | -20.49 | -24.84 | -4.36 | 3.62 |
| 56 | E9PR52 | Chitinase-3-like protein 2 | CHI3L2 | -21.29 | -25.72 | -4.43 | 3.6 |
| 57 | P02788-2 | Lactotransferrin | LTF | -18.8 | -24.04 | -5.24 | 3.38 |
| 58 | P18206-2 | Vinculin | VCL | -22.71 | -26.28 | -3.58 | 3.34 |
| 59 | P52566 | Rho GDP-dissociation inhibitor 2 | ARHGDIB | -18.87 | -22.34 | -3.47 | 3.31 |
| 60 | P62942 | Peptidyl-prolyl cis-trans isomerase FKBP1A | FKBP1A | -19.83 | -24.1 | -4.27 | 3.27 |
| TBM group |  |  |  |  |  |  |  |
| No | Protein ID | Protein name | Gene name | Mean intensity of BM (log<sub>2</sub>) | Mean intensity of Other (log<sub>2</sub>) | Difference in intensity between TBM and Others | -Log (p value) |
| 1 | P25311 | Zinc-alpha-2-glycoprotein | AZGP1 | -16.21 | -17.29 | -1.08 | 9.22 |
| 2 | P23381 | Tryptophan-tRNA ligase | WARS | -20.06 | -23.81 | -3.75 | 5.58 |
| 3 | P29622 | Kallistatin | SERPINA4 | -20.65 | -22.77 | -2.13 | 4.39 |
| 4 | P02746 | Complement C1q subcomponent subunit B | C1QB | -18.32 | -19.31 | -0.99 | 4.35 |
| 5 | A0A075B6J0 | Immunoglobulin lambda variable 1-40 | IGLV1-40 | -17.36 | -20.25 | -2.9 | 4.25 |
| 6 | P02749 | Beta-2-glycoprotein 1 | APOH | -18.32 | -19.33 | -1.01 | 4.13 |
| 7 | P32455 | Guanylate-binding protein 1 | GBP1 | -23.23 | -25.44 | -2.21 | 3.41 |
| 8 | P16070-10 | CD44 antigen | CD44 | -22.26 | -23.46 | -1.19 | 3.23 |
| 9 | P02747 | Complement C1q subcomponent subunit C | C1QC | -17.8 | -18.93 | -1.13 | 3.14 |
| 10 | P01591 | Immunoglobulin J chain | JCHAIN | -19.12 | -22.1 | -2.98 | 2.93 |
| 11 | Q8WVN6 | Secreted and transmembrane protein 1 | SECTM1 | -20.96 | -23.77 | -2.81 | 2.88 |
| 12 | Q96IY4 | Carboxypeptidase B2 | CPB2 | -21.79 | -24.05 | -2.26 | 2.83 |
| 13 | Q14624 | Inter-alpha-trypsin inhibitor heavy chain H4 | ITIH4 | -22.42 | -24.14 | -1.72 | 2.82 |
| 14 | P01625 | Immunoglobulin kappa variable 4-1 | IGKV4-1 | -23.58 | -26.87 | -3.29 | 2.78 |
| 15 | O15204 | ADAM DEC1 | ADAMDEC1 | -22.75 | -25.26 | -2.51 | 2.64 |
| 16 | P19971-2 | Thymidine phosphorylase | TYMP | -22.78 | -25.01 | -2.24 | 2.57 |
| 17 | A0A075B6J9 | Immunoglobulin lambda variable 2-18 | IGLV2-18 | -21.35 | -24.22 | -2.87 | 2.37 |
| 18 | P01596 | Immunoglobulin kappa variable 1-5 | IGKV1-5 | -21.02 | -23.7 | -2.69 | 2.36 |
| 19 | P02743 | Serum amyloid P-component | APCS | -23.62 | -25.81 | -2.19 | 2.3 |

**Table S4.** Results of analysis comparing the diagnostic value of LCN2 in distinguishing between patients with confirmed and clinically suspected bacterial meningitis.

| **cBM vs. sBM** | **Cut-off** | **AUC**<br>**(95% CI)** | **Sensitivity**<br>**(95% CI)** | **Specificity**<br>**(95% CI)** | **DOR**<br>**(95% CI)** |
| --- | --- | --- | --- | --- | --- |
| LCN2 (ng/ml) | 452.3 | 0.74<br>(0.6-0.88) | 0.82<br>(0.68-0.9) | 0.6<br>(0.39-0.78) | 6.8<br>(2.1-21.9) |
| CSF leukocytes (cell per mm<sup>3</sup>) | 1533 | 0.61<br>(0.47-0.76) | 0.61<br>(0.47-0.74) | 0.65<br>(0.43-0.82) | 3.0<br>(1-8.9) |
| CSF lactate (mmol/L) | 9.0 | 0.84<br>(0.74-0.95) | 0.77<br>(0.63-0.87) | 0.8<br>(0.58-0.92) | 13.6<br>(3.7-50.1) |
| CSF/blood glucose ratio | 0.2 | 0.76<br>(0.63-0.88) | 0.57<br>(0.42-0.7) | 0.95<br>(0.76-0.99) | 25<br>(3.1-203.6) |
| CSF protein (g/L) | 2.1 | 0.74<br>(0.61-0.86) | 0.66<br>(0.51-0.78) | 0.75<br>(0.53-0.89) | 5.8<br>(1.8-19) |
| CSF White cell count+lactate+CSF/blood glucose level+CSF protein | NA | 0.87<br>(0.77-0.97) | 0.95<br>(0.85-0.99) | 0.65<br>(0.43-0.82) | 39<br>(7.2-21.4) |
| CSF White cell count+lactate+CSF/blood glucose level+CSF protein+lipocalin 2 | NA | 0.87<br>(0.77-0.96) | 0.95<br>(0.85-0.99) | 0.65<br>(0.43-0.82) | 39<br>(7.2-21.4) |

**Figure S1.**
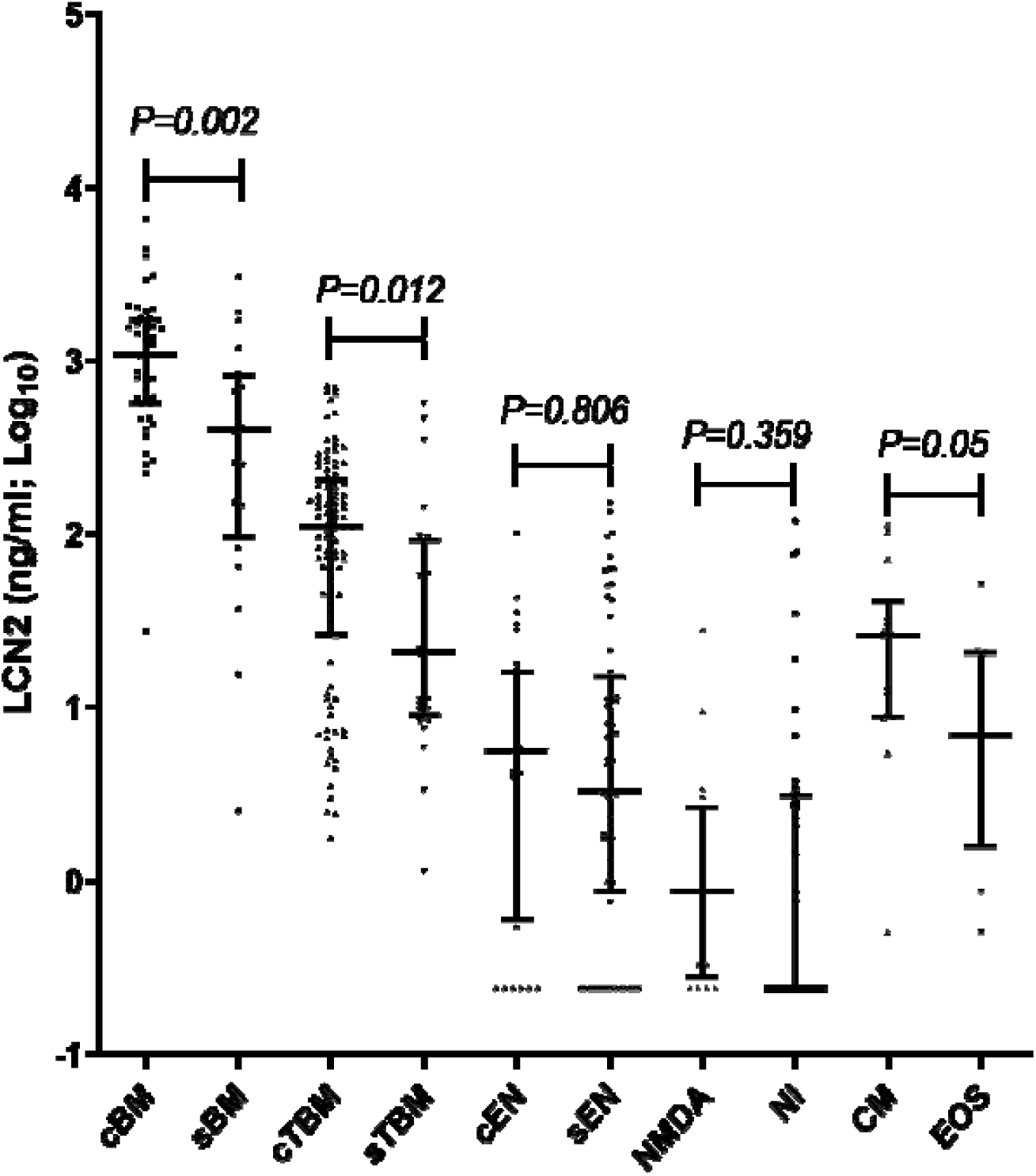
Plots showing the distribution of LCN2 concentrations in patients with laboratory confirmed or clinically suspected CNS infections and non-CNS infections. **Note to Figure S1:** cBM: confirmed bacterial meningitis, sBM: clinically suspected bacterial meningitis, cTBM: confirmed tuberculous meningitis, sTBM: clinically suspected tuberculous meningitis, cEN: confirmed encephalitis, sEN: clinically suspected encephalitis, NMDA: anti-NDMAR encephalitis, NI: non-CNS infections, CM: crytococcal meningitis, EOS: eosinophilic meningitis

**Figure S2.**
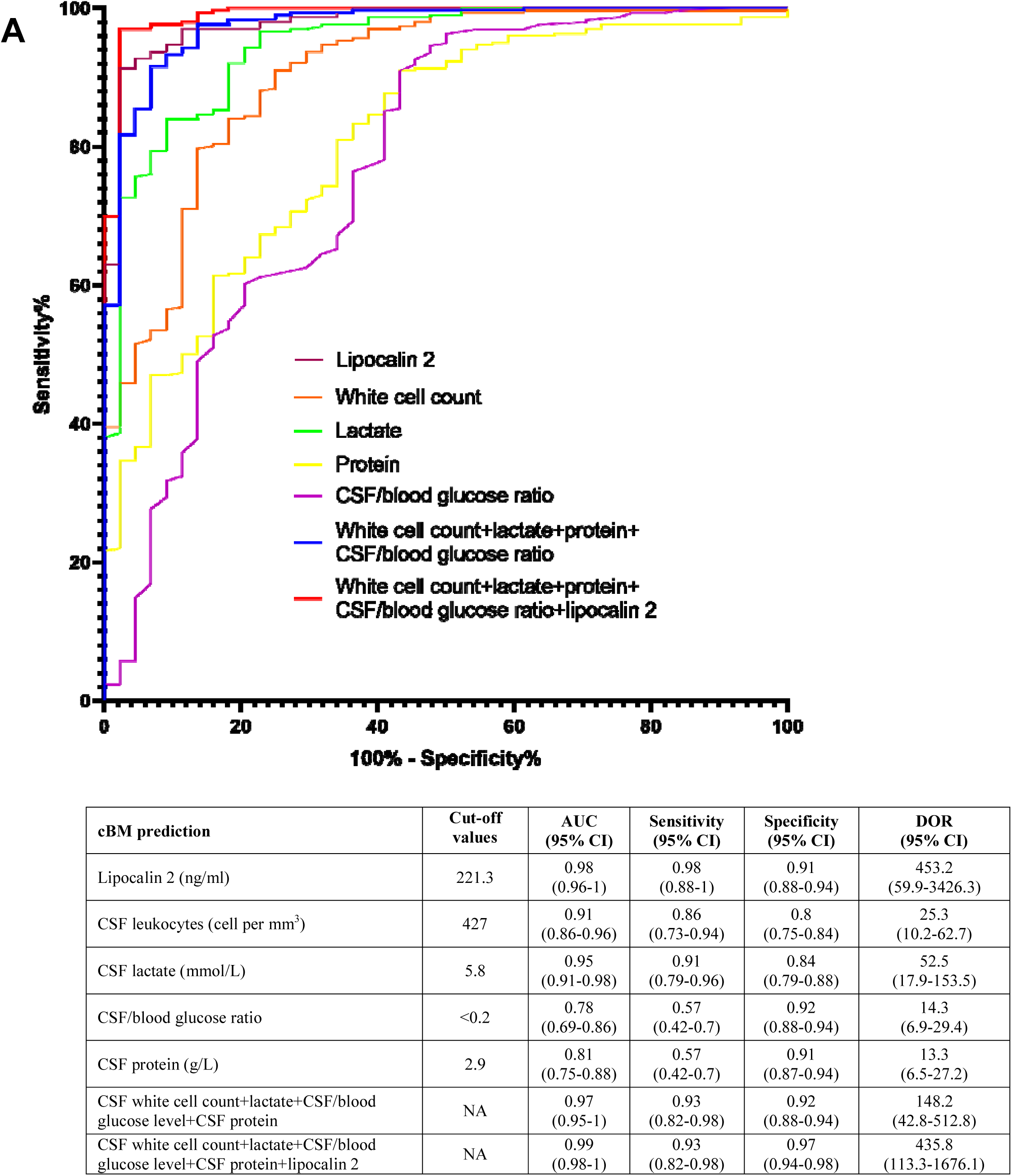
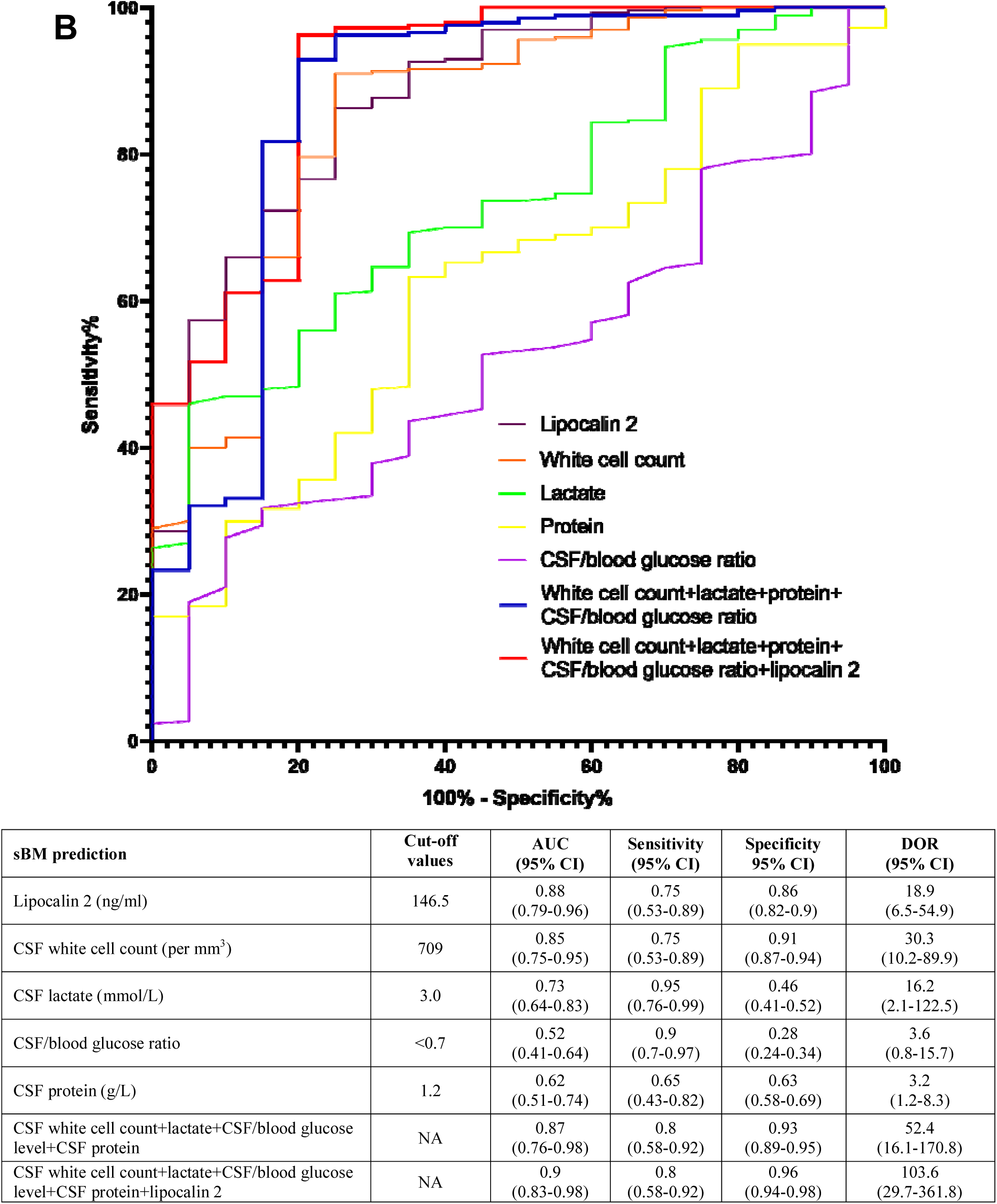
Diagnostic performance of LCN2 in discriminating between laboratory confirmed (A) or clinically suspected bacterial meningitis patients (B) and other clinical entities and in comparison with existing biomarkers. **Note to Figure S2:** NA: not applicable

